# Human iPSC-derived motor neuron innervation enhances the differentiation of muscle bundles engineered with benchtop fabrication techniques

**DOI:** 10.1101/2024.12.02.626391

**Authors:** Jeffrey W. Santoso, Stephanie K. Do, Riya Verma, Alexander V. Do, Eric Hendricks, Justin K. Ichida, Megan L. McCain

## Abstract

Engineered skeletal muscle tissues are critical tools for disease modeling, drug screening, and regenerative medicine, but are limited by insufficient maturation. Because innervation is a critical regulator of skeletal muscle development and regeneration in vivo, motor neurons are hypothesized to improve the maturity of engineered skeletal muscle tissues. Although motor neurons have been added to pre-engineered muscle constructs, the impact of motor neurons added prior to the onset of muscle differentiation has not been evaluated. In this study, benchtop fabrication equipment was used to facilely fabricate chambers for engineering 3-dimensional (3-D) skeletal muscles bundles and measuring their contractile performance. Primary chick myoblasts were embedded in an extracellular matrix hydrogel solution and differentiated into engineered muscle bundles, with or without the addition of human induced pluripotent stem cell (hiPSC)-derived motor neurons. Muscle bundles differentiated with motor neurons had neurites distributed throughout their volume and a higher myogenic index compared to muscle bundles without motor neurons. Innervated muscle bundles also generated significantly higher twitch and tetanus forces in response to electrical field stimulation after one and two weeks of differentiation compared to non-innervated muscle bundles cultured with or without neurotrophic factors. Non-innervated muscle bundles also experienced a decline in rise and fall times as the culture progressed, whereas innervated muscle bundles and non-innervated muscle bundles with neurotrophic factors maintained more consistent rise and fall times. Innervated muscle bundles also expressed the highest levels of the genes for slow myosin light chain 3 (*MYL3*) and myoglobin (*MB*), which are associated with slow twitch fibers. These data suggest that motor neuron innervation enhances the structural and functional development of engineered skeletal muscle constructs and maintains them in a more oxidative phenotype.

## Introduction

Throughout all stages of life, the maturity of skeletal muscle fibers is highly regulated by motor neuron input. During embryonic development, muscle progenitors autonomously fuse into primary myotubes, but require innervation to form secondary myotubes, which generate most adult muscle fibers^1, 2^. In adult rat, denervated muscle undergoes atrophy and functional decline^3^ and regenerates to a lesser extent after injury^4^. However, denervated muscle can be recovered by reinnervating it with transplanted nerve-muscle-endplate bands^3^. Age-related muscular atrophy^5^ and muscle wasting in many neuromuscular disorders, such as spinal muscular atrophy^6, 7^, has also been attributed to motor neuron degeneration and denervation. Interestingly, neurotrophic factors alone, such as brain-derived neurotrophic factor (BDNF), ciliary neurotrophic factor (CNTF), and glial cell-derived neurotrophic factor (GDNF), have also been shown to rescue denervation-induced muscle atrophy^8, 9^, demonstrating the functional importance of neuron-associated paracrine signaling on muscle growth.

Engineered skeletal muscle constructs are critically needed as in vitro disease models and as grafts that could be transplanted into patients. However, the maturity of engineered muscle tissues remains limited. Because innervation is a key driver of muscle maturation in vivo, innervation may also promote maturation of engineered muscle tissues in vitro. In previous studies, engineered muscle tissues have been innervated primarily by adding motor neurons to muscle constructs after they are differentiated into myotubes in various 2-dimensional (2-D)^10–13^ or 3-dimensional (3-D) configurations^14–18^. In these settings, motor neurons increased the expression of genes important for the contractile machinery but had no significant impact on myotube formation^19^ or the amplitude of force generated by the muscle in response to electrical stimulation^17^. Thus, innervation generally had minimal impact on muscle maturation when it was introduced post-myotube differentiation. The effect of innervation introduced at earlier stages of muscle differentiation has not been investigated.

Given the importance of innervation in the early development of endogenous skeletal muscle, we hypothesized that skeletal myoblasts would generate more mature muscle constructs if they were differentiated into myotubes in the presence of motor neurons. To test this, we first developed a method for benchtop fabrication of 3-D muscle bundles and optical measurements of force generation. We then engineered innervated and non-innervated muscle bundles from primary chick myoblasts with and without human induced pluripotent stem (hiPSC) cell-derived motor neurons, respectively. To evaluate the structure and function of the resulting constructs, we performed quantitative immunostaining, measured twitch and tetanus forces, and quantified the expression of a panel of fast and slow muscle fiber genes. To determine any impacts of neurotrophic factors on muscle differentiation, we also cultured non-innervated muscle bundles with or without neurotrophic factors. Our results indicate that innervated muscle bundles had enhanced structural and functional maturation compared to their non-innervated counterparts and expressed higher levels of the myoglobin gene, suggestive of a more oxidative phenotype. Neurotrophic factors also induced some improvements in muscle phenotype, but to a lesser extent than motor neurons. Thus, introducing motor neurons during muscle differentiation significantly improved the structural and functional maturation of engineered muscle tissues, which has important implications for understanding mechanisms of muscle development and improving the maturity of engineered muscle tissues for in vitro modeling and transplantation.

## Methods

### Fabrication of 3D Muscle Bundle Devices

Based on the dimensions established in our previous protocol^20^, muscle bundle templates comprising two 13 x 2 x 1 (length x width x height) mm half-chambers were designed in TinkerCAD (Autodesk Inc.). Towards the ends of each half-chamber, the width was flared to 3.5 mm to improve visualization of the PDMS anchor rods for downstream analysis (**Supplemental Figure 1A-C**). The design was 3-D printed in resin using a CADWorks3D M50 Series digital light processing printer (CADWorks3D). Sylgard 184 PDMS was cured within the templates overnight at 65°C and carefully removed using the circular end tab and a flat spoon spatula.

To fabricate the anchor rod layer, a 30W Epilog Mini 24 Laser Engraver (70% speed, 45% power, 5000 Hz frequency) was used to engrave 500 μm-wide and 2 mm-long rods from a 0.25 mm-thick commercial silicone sheet (Rogers HT-6240-0.25) (**Supplemental Figure 1D**). The cut sheet was then plasma treated for ten minutes (Harrick Plasma, PDC-001-HP) and sandwiched between the two PDMS half-chambers.

Devices were reversibly bonded onto the bottom well surfaces of a 6-well plate by treating both surfaces for 8 minutes in a UVO Cleaner (Jelight Company Model 342). Devices were stored at ambient conditions until the day prior to cell seeding, at which point they were incubated overnight in a biosafety cabinet under UV light for sterilization.

### Isolation of Primary Chick Myoblasts

Thigh muscle tissue from day 10 chick embryos (AA Lab Eggs) was isolated and dissociated into myoblasts as previously described^12, 21^. Briefly, minced muscle was digested with 1 mg/mL collagenase (Worthington LS004177, Lot 43K144303B). The resulting cell solution underwent two 30-minute pre-plating steps to purify myoblasts. Myoblasts were either expanded in T175 flasks in growth medium containing 10% horse serum and 5% chicken serum or immediately used for experiments. Cells cultured up to 3 passages were used for experiments. Day 10 chick embryos are not considered live vertebrate animals and thus not subject to IACUC review.

### Differentiation and culture of hiPSC-derived motor neurons

Human induced pluripotent stem cell (hiPSC)-derived motor neurons were generated as previously described^12, 22^. Briefly, lymphocytes attained from the NINDS Biorepository at the Coriell Institute for Medical Research (ND03231) were previously reprogrammed into hiPSCs with the use of episomal vectors containing Oct4, Sox2, Klf4, c-Myc, Lin28, and a p53 shRNA^22,23^. hiPSCs were expanded and maintained in mTeSR media until differentiation. Then, a small molecule cocktail that included CHIR99021, DMH1, and SB431542 was incorporated into the differentiation media formulation described in the previous section without ACA to generate OLIG2-positive motor neuron progenitors. Accutase (Gibco A1110501) was used to detach progenitors from the culture surface, which were then re-seeded onto untreated, low-attachment polystyrene dishes to induce formation of spheroidal aggregates. Subsequent enrichment of the progenitors into functional motor neurons (>90%) occurred over another 16 days with the use of a Notch inhibitor (compound E) and activation of Hedgehog signaling (purmorphamine)^22, 24^.

### Engineering Innervated and Non-Innervated Muscle Bundles

The muscle bundle chambers of sterilized devices were treated with 1% w/v Pluronic solution (diluted from 20% with phosphate buffered saline, Invitrogen P3000MP) for one hour prior to seeding chick myoblasts. Then, 1.5 million chick myoblasts were suspended in 32.5 μL of growth media and 4 μL of thrombin (50 U/mL in sterile water, Sigma-Aldrich T4648). Another vial of 20 μL growth media, 20 μL fibrinogen (20 mg/mL in growth media, Sigma-Aldrich F8630), and 20 μL Matrigel hESC-Qualified Matrix (Corning 354277) was mixed and kept on ice^25, 26^. The myoblast-thrombin solution was then mixed rapidly, taking care to not form bubbles, with the fibrinogen solution on ice. 70 μL of solution (∼1 million cells) was pipetted into the chamber of each device, gelling around the PDMS anchor rod layer within 30 seconds. The hydrogel was then allowed to solidify at 37 °C for 30 minutes before growth media supplemented with 1.5 mg/mL of aminocaproic acid (ACA, Sigma Aldrich), an anti-fibrinolytic agent, was added to fully submerge the muscle bundles.

After four days of culture in growth media supplemented with ACA, muscle bundle devices were detached from the bottoms of the 6-well plate and cultured on a rocker (0.4 Hz) in serum-free differentiation medium containing ACA, N-2, and B-27 supplements (Gibco, 17502-048 and 17504-044) to increase nutrient diffusion^27^. Media was exchanged every other day.

To generate innervated chick muscle bundles, five motor neuron spheroids were isolated manually and partially dissociated by pipetting 20 times through a P1000 and P20 pipette tip, in succession. Then, the partially dissociated motor neurons were added to the chick myoblast and thrombin solution detailed in the previous section prior to mixing with the fibrinogen solution. Following addition of the mixture into the device, the fibrin hydrogel was allowed to solidify at 37 °C for 30 minutes, and growth media with the addition of 1.5 mg/mL ACA and 10 ng/mL of brain-derived neurotrophic factor (BDNF), ciliary neurotrophic factor (CNTF), and glial cell derived neurotrophic factor (GDNF) were added to submerge bundles. Media was exchanged after 2 days, and after 4 days in growth media with ACA and neurotrophic factors, media was switched to differentiation medium with ACA and neurotrophic factors and cultured on a rocker, as described above. Media was then exchanged every other day, with neurotrophic factors removed after one week of differentiation. Select muscle bundles without motor neurons were also cultured in media with neurotrophic factors for the same time period as those with motor neurons.

### Structural Characterization

After one or two weeks of differentiation, muscle bundles were rinsed in phosphate buffered saline (PBS) and fixed with ice-cold methanol for 30 minutes at 4°C. After three PBS rinses, muscle bundles were incubated in 30% sucrose overnight at 4°C to displace water and prevent ice crystal formation during freezing^20^. Samples were then immersed in optimal cutting temperature compound (Sakura Finetek 4583) in 25 x 20 x 5 mm cryomolds and incubated at room temperature for 30 minutes. Then, samples were frozen in liquid nitrogen-chilled isopentane for 1 minute. Frozen myobundles were kept at -80°C until sectioning. A Leica CM3050 S Research Cryostat was used to generate 12 μm thick longitudinal sections.

Select sections were stained for hematoxylin and eosin (H&E) following standard protocols. Other sections were incubated in 10% goat serum for 1 hour and then primary antibodies for mouse sarcomeric α-actinin (Sigma, A7811) and rabbit synapsin-1 (Cell Signaling Technology, D12G5), both at a 1:200 dilution. Secondary staining was performed using goat anti-rabbit antibody conjugated to Alexa Fluor 488, goat anti-mouse antibody conjugated to Alexa Fluor 546, and DAPI (all 1:200). Confocal microscopy was performed using a Nikon Eclipse Ti microscope at 20X air objective with Confocal Module Nikon C2 and Andor Zyla sCMOS camera.

ImageJ (NIH) was used to analyze images of longitudinal sections. Number of nuclei per area was quantified by counting the number of nuclei, as determined by DAPI thresholding, and dividing by the area of the myobundle, as determined by the area inside the boundaries of the α-actinin stain, per field of view. The myogenic index, which represents the proportion of nuclei incorporated into α-actinin positive myotubes, was quantified by dividing the number of nuclei colocalized in thresholded α-actinin stains with the total number of nuclei. For innervated myobundles, synapsin stain was thresholded to determine the presynapse area. Nuclei in these areas were excluded from nuclei density and myogenic index analysis. Lastly, bundle width was determined by measuring the maximum distance between the top and bottom of bundles as marked by the H&E stain.

### Quantifying Contractile Force

After one or two weeks of differentiation, muscle bundles were transferred to a 35 mm Petri dish, placed on the stage of a Nikon SMZ745T stereomicroscope, and incubated in standard Tyrode’s solution, similar to previous protocols^12^. Platinum field electrodes were inserted into the bundle chamber, perpendicular to the muscle bundle. Twitch and tetanus contractions were stimulated by a 30 V/cm electrical stimulus at 2 and 20 Hz, respectively. Ten-second videos were recorded at 100 frames per second with a Basler acA640-120um USB 3.0 camera.

ImageJ tools were used to manually trace the angle of deflection (θ) of PDMS rods anchoring both sides of the muscle bundle. This angle was used to calculate displacement of the PDMS rods, which was then converted to force by referencing a linear standard curve, which was created by applying known weights to PDMS rods and measuring the angle of deflection. Time dynamics of contractions were analyzed by selecting a rectangular region of interest on the PDMS rods and tracking changes in the total pixel intensity of the region over time. Rise and fall times were calculated using the risetime and falltime function in MATLAB between the 10% and 90% reference level of the waveform.

### Quantifying Gene Expression with RT-qPCR

After one or two weeks of differentiation, bundles were removed from their PDMS mold and roughly minced with a sterile X-acto knife before being lysed in 1 ml of TRIzol reagent (Thermo Fisher Scientific) for 30 minutes on ice. mRNA was extracted through the aqueous layer of a TRIzol-chloroform phase separation, and the RNA isolation was performed using a miRNeasy kit and protocol (Qiagen). Reverse transcription was used to synthesize cDNA using iScript Reverse Transcription Supermix for RT-qPCR kit (Bio-Rad Laboratories). Primers for PCR were designed using the National Center for Biotechnology Information (NCBI) Primer Design Tool. All primer information is listed in **Supplemental Table 1.** RT-qPCR was conducted using the SsoAdvanced Universal SYBR Green Supermix (Bio-Rad Laboratories) with the CFX384 Touch Real-Time PCR Detection System (Bio-Rad Laboratories). The generated cycle threshold (Ct) values for each target gene were normalized relative to the housekeeping gene *ACTB*. The fold change for each gene was calculated using 2−ΔΔCt^28^.

### Statistical Analysis

All data were tested for normality and analyzed using student’s t-test, Mann-Whitney test, one-way ANOVA, or two-way ANOVA as appropriate, using GraphPad Prism 8. Comparisons with p-values less than 0.05 were considered statistically significant. In figures, * denotes p<0.05, ** p<0.01, *** p<0.001, **** p<0.0001, unless otherwise noted.

## Results

### Benchtop Fabrication of Engineered Muscle Bundle Templates

We first developed a method to fabricate a device for engineering elongated muscle bundles with an integrated system for measuring force generation via optical imaging using only benchtop equipment. Each device consists of three bonded layers of PDMS (**Figure 1A)**. The top and bottom layers are 1 mm thick slabs with 13 mm x 2 mm rectangular voids replica molded from 3-D printed templates (**Figure 1B)**, similar to our previous approach^20^. These pieces form the chamber that confines the muscle cells and media. The middle layer is a PDMS sheet laser-cut with a similar rectangular void and two thin PDMS rectangles (or rods) that extend across the width of the rectangle at the longitudinal ends (**Figure 1C**). The thin PDMS rods are designed to anchor the muscle bundle and deflect with contraction for visual tracking. To improve the visualization of PDMS rod deflection, we flared the width of the chamber at the longitudinal ends. To enable force measurements based on PDMS rod deflection, we hung a series of weights on PDMS rods and measured the corresponding displacement to generate a displacement-force calibration curve (**Figure 1D**).

**Figure 1:**
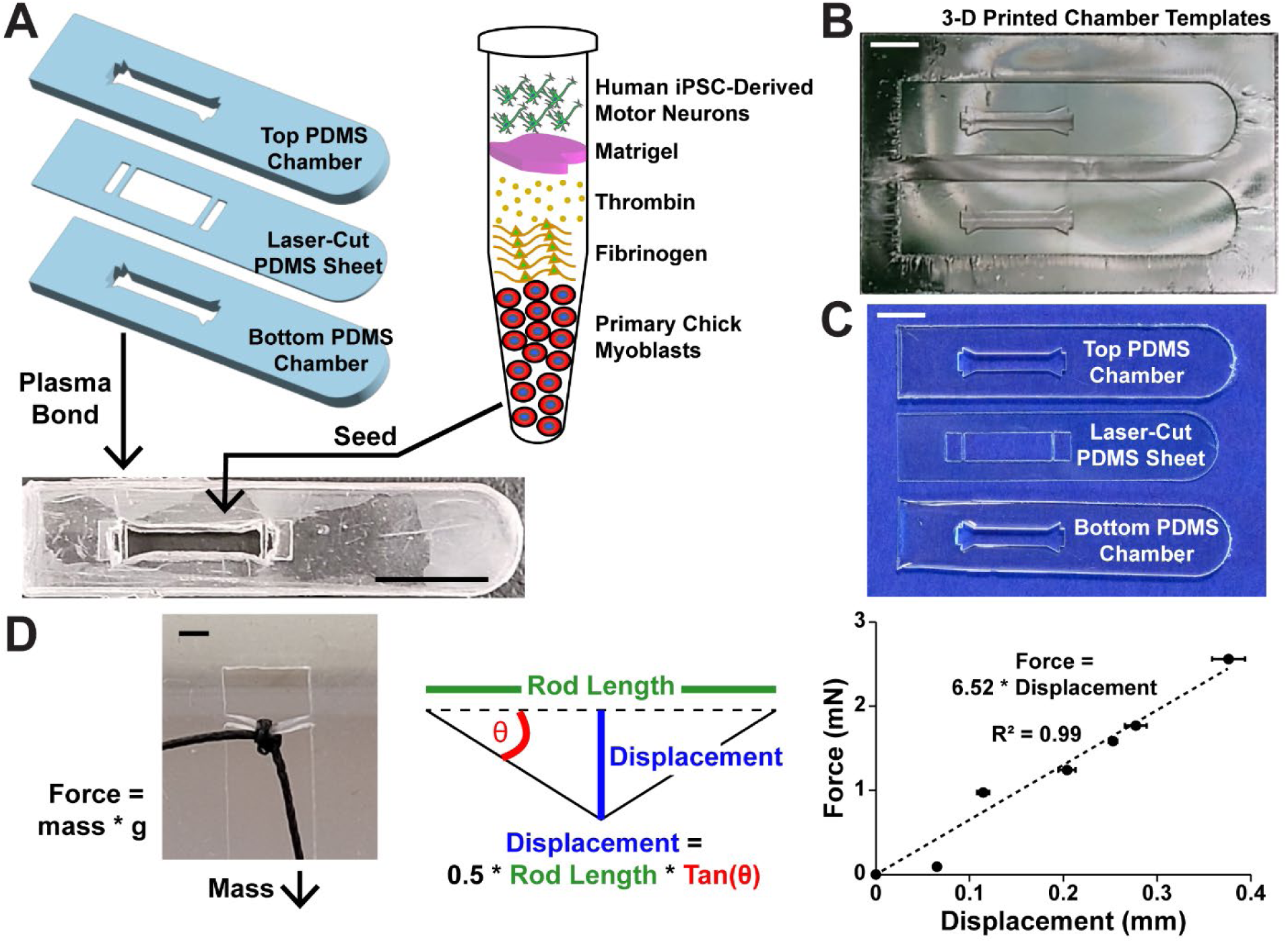
Fabrication and calibration of muscle bundle devices. (A) Photograph of 3-D printed template for top and bottom pieces of muscle bundle chamber. Scale bar: 5 mm. (B) Photograph of top and bottom PDMS components molded from 3-D printed templates. (C) Photograph of top and bottom PDMS components and laser-cut PDMS sheet containing the rods to anchor muscle bundles. Scale bar: 5 mm. (D) Photograph (left) and calculation (middle) used to generate force-displacement relationship (right) to calibrate rod displacement to muscle bundle force generation. Scale bar: 1 mm.

To engineer muscle bundles, we bonded the 3-layered PDMS device to the bottom of a 6-well plate well and injected a solution of fibrin hydrogel pre-polymer and primary chick skeletal myoblasts, similar to previous protocols^25, 26^. To engineer innervated muscle bundles, we added clusters of human iPSC-derived motor neurons to the solution. For both types of bundles, cells were cultured for four days in growth media, then switched to differentiation media and cultured for an additional one or two weeks on a rocking platform. Innervated muscle bundles were also cultured with neurotrophic factors BDNF, CNTF, and GDNF to support motor neuron viability for the first week of culture. As expected, the muscle cells compacted around the PDMS rods and formed aligned bundles with consistent width over the two-week differentiation period, as evaluated by sections stained with H&E (**Figure 2A-B)**. Muscle bundles with hiPSC-derived motor neurons were less compact after one week, likely due to steric hindrance from the neuron clusters, but eventually compacted to similar levels as the non-innervated bundles (**Figure 2A-B)**. Localized clusters of motor neurons were still visible at two weeks, but these did not prevent the muscle cells from ultimately compacting into an aligned bundle.

**Figure 2:**
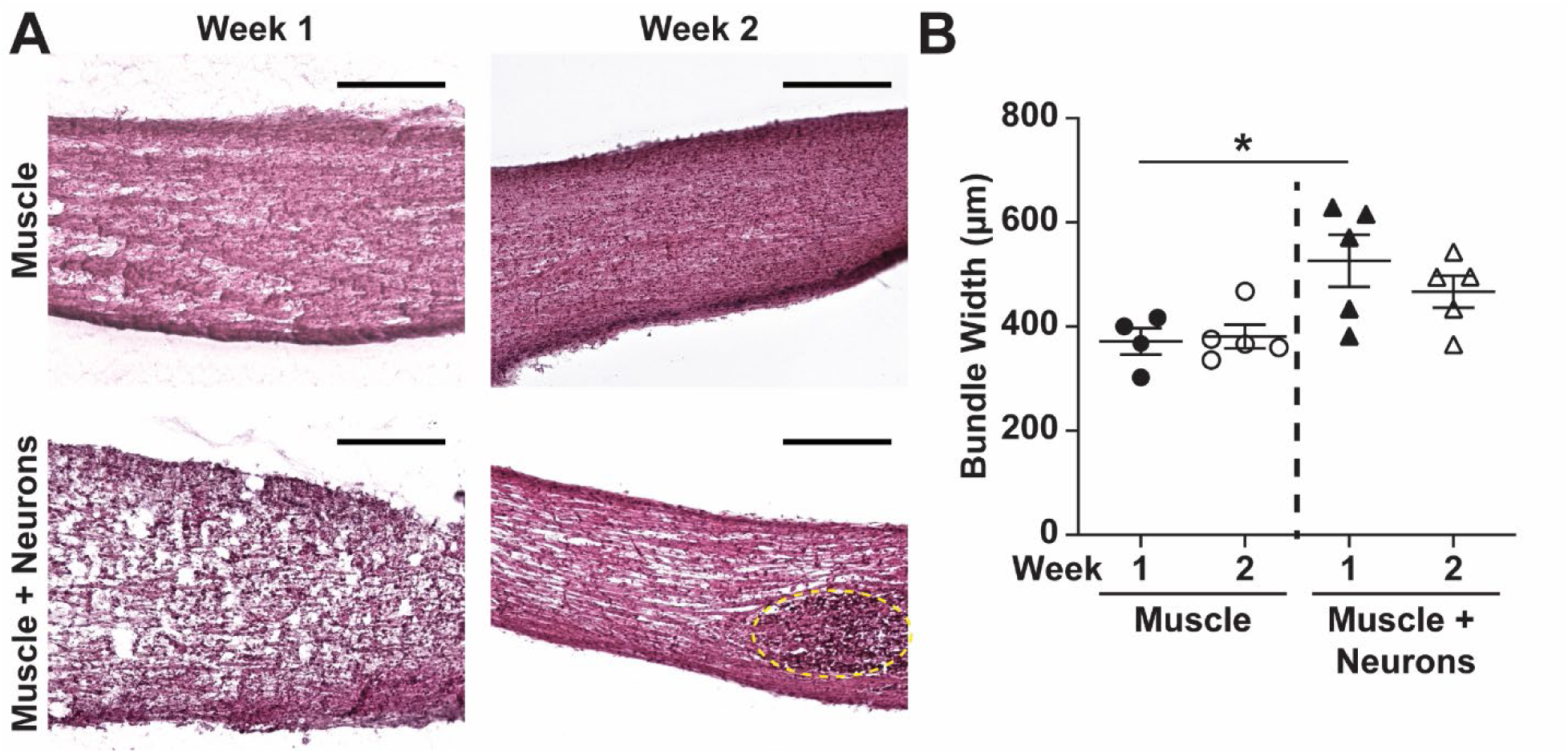
Histology of non-innervated and innervated muscle bundles. (A) Hematoxylin and eosin stained muscle bundle sections differentiated with or without hiPSC-derived motor neurons for one or two weeks. Yellow dotted circle marks a cluster of neurons. Scale bar: 200 μm. (B) Bundle width at weekly timepoints for muscle bundles with and without neurons.

### Impact of Innervation on Muscle Bundle Structure

To closely evaluate cell and tissue architecture, we next stained sections of non-innervated and innervated muscle bundles after one and two weeks of differentiation with DAPI and sarcomeric alpha-actinin (**Figure 3A)**. At two weeks, both non-innervated and innervated bundles comprised aligned muscle fibers with well-formed sarcomeres. The number of nuclei per area was slightly lower in innervated bundles compared to non-innervated bundles at one week (**Figure 3B)**, consistent with the lower degree of compaction observed in the H&E staining (**Figure 2A)**. However, this difference was not statistically significant and the number of nuclei at week two was more similar between non-innervated and innervated bundles (**Figure 3B**). The myogenic index was significantly higher in innervated bundles compared to non-innervated bundles at both timepoints (**Figure 3C)**, suggestive of more muscle differentiation with innervation. To evaluate motor neuron morphology, we also stained innervated muscle bundles for the pre-synaptic marker synapsin and quantified the area of synapsin coverage, which significantly increased from one to two weeks (**Figure 3D)**. This suggests that motor neurons were mostly confined to their initial clusters after one week, but gradually migrated and extended axons throughout the muscle fibers as the culture time progressed, as shown in **Figure 3A**. However, due to technical challenges related to sectioning and immunostaining of the engineered 3-D constructs, we were unable to resolve individual neuromuscular junctions and thus cannot confirm formation of synapses.

**Figure 3:**
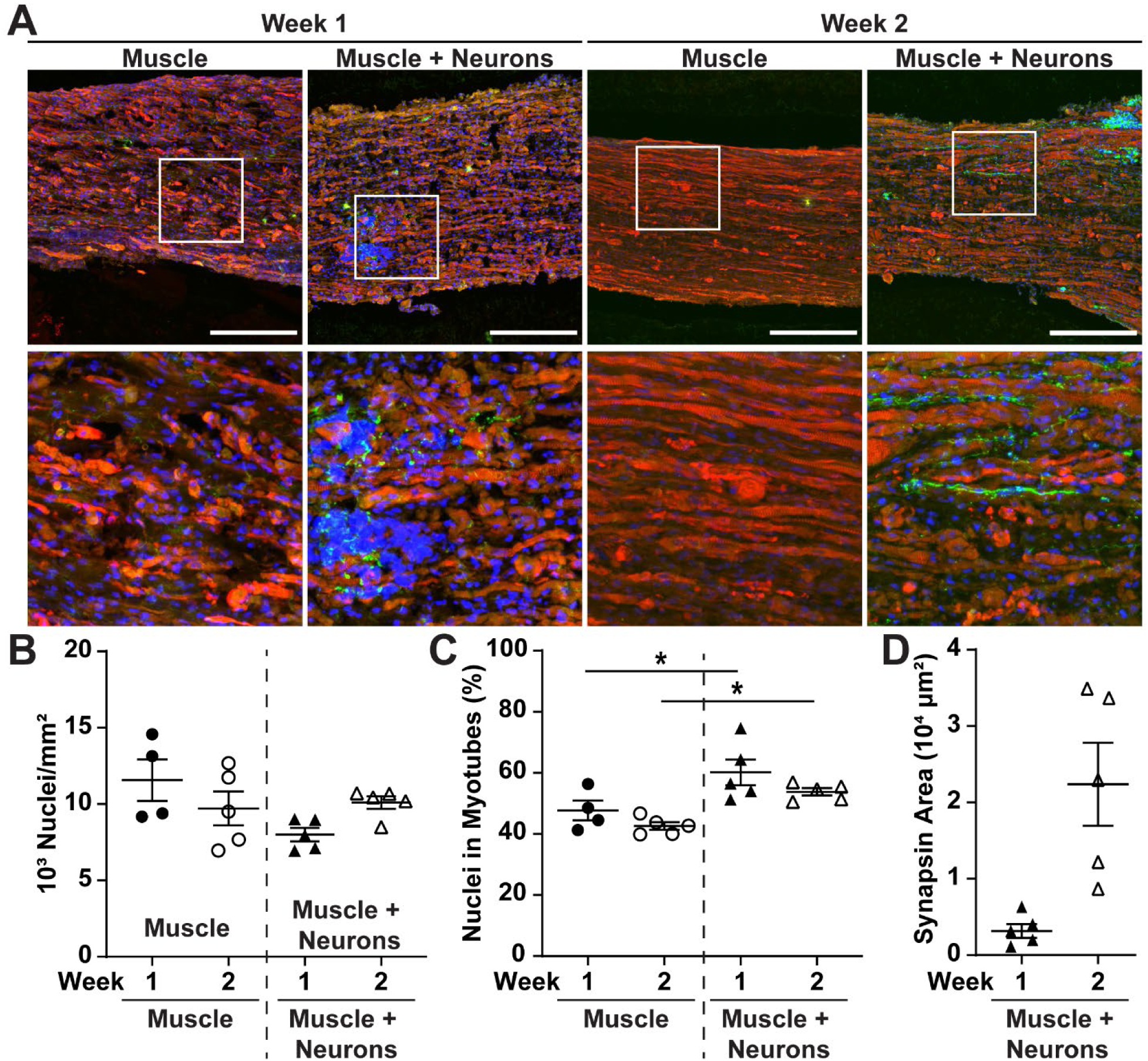
Muscle differentiation in non-innervated and innervated muscle bundles. (A) Muscle bundle sections stained with DAPI (blue) and sarcomeric alpha-actinin (red) differentiated with or without hiPSC-derived motor neurons for one or two weeks. Bundles with neurons were also stained for synapsin (green). In top row, scale bar: 200 μm. In bottom row, scale bar: 50 μm. (B) Number of nuclei normalized to area, (C) myogenic index, and (D) synapsin area at weekly timepoints for muscle bundles with and without neurons.

### Impact of Innervation and Neurotrophic Factors on Muscle Bundle Contractility

Our next goal was to evaluate the impact of hiPSC-derived motor neurons on the contractile output of engineered muscle tissues. As mentioned above, for the first week of culture, our innervated muscle bundles were cultured with the neurotropic factors BDNF, CTNF, and GDNF to support motor neuron viability. However, these neurotropic factors have been previously demonstrated to induce myotropic effects on their own, including elevating formation of fast twitch fibers, improving denervation-induced decline in contractile forces, and shortening the time of contraction^9, 29, 30^. Thus, we also cultured non-innervated muscle bundles with neurotropic factors for the same amount of time (first week of culture) as innervated muscle bundles. After one and two weeks of culture, we stimulated twitch or tetanus forces by pacing the engineered bundles with a field stimulation electrode at 2 Hz and 20 Hz, respectively. We tracked the deflection of the PDMS rods (**Figure 4A**) and converted maximum displacement to force using our calibration curve (**Figure 1D**). For each batch of engineered tissues, we then normalized all force measurements to the average value for non-innervated muscle bundles without neurotrophic factors to account for variability in the primary harvest of chick myoblasts. As shown in **Figure 4B** and **4C**, normalized twitch forces and tetanus forces were significantly higher in innervated muscle bundles (**Supplemental Movie 1**) compared to non-innervated muscle bundles (**Supplemental Movie 2**) at both one- and two-week timepoints. The presence of neurotrophic factors had no impact on the normalized magnitude of contraction force (**Figure 4B, C).** The tetanus-to-twitch ratio, which corresponds to muscle maturity, was higher in innervated muscle bundles after one week post differentiation, but was not statistically different than non-innervated muscle bundles at two weeks post differentiation (**Figure 4D).**

**Figure 4:**
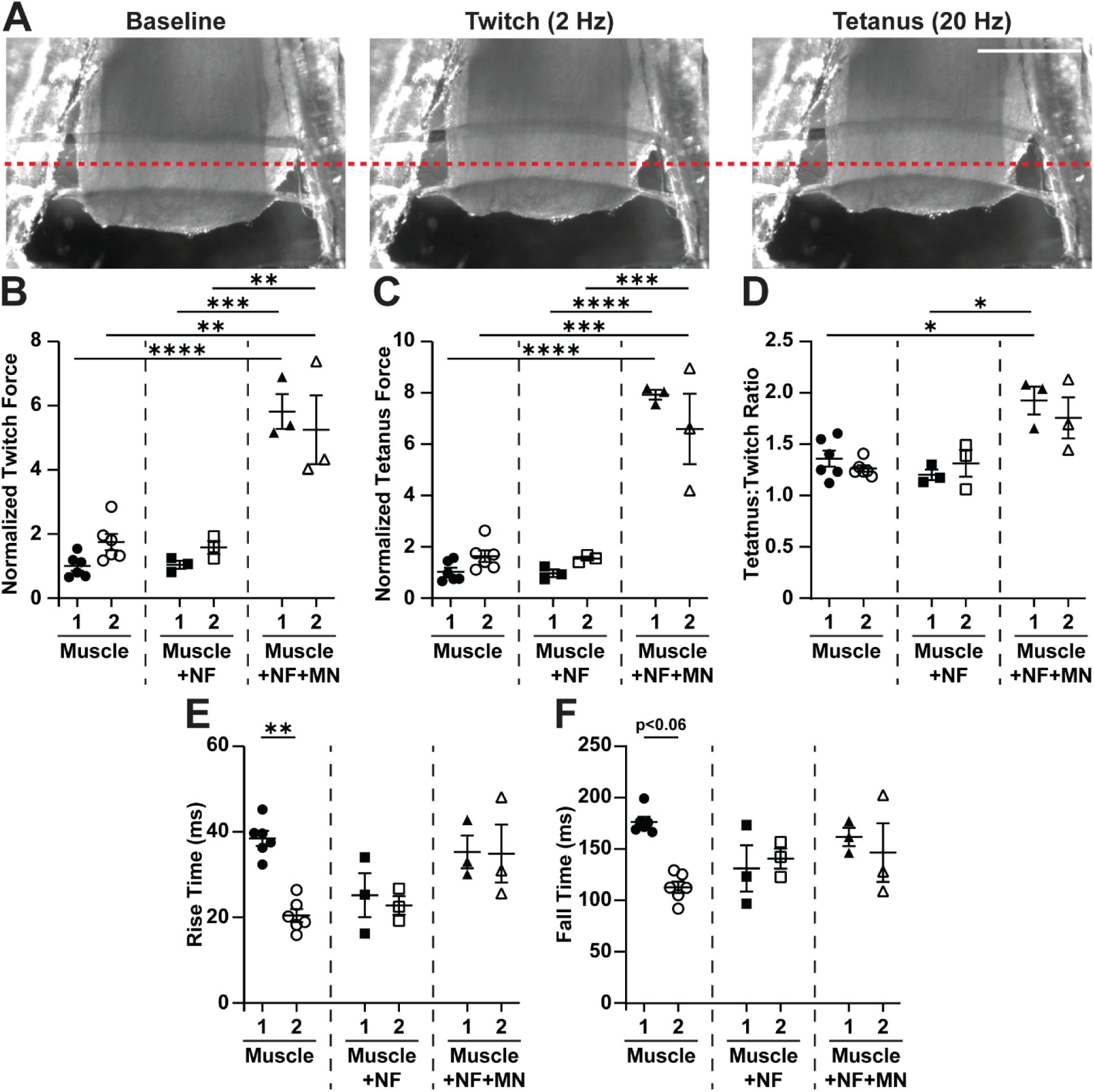
Electrically stimulated force generation in muscle bundles cultured with or without neurotrophic factors (NF) or motor neurons (MN). (A) Frames of an innervated muscle bundle at baseline, peak twitch, and peak tetanus. Red dotted lines are for reference. Scale bar: 1 mm. Electrically stimulated (B) twitch force, (C) tetanus force, (D) tetanus-to-twitch ratio, (D) twitch rise time, and (E) twitch fall time for each type of muscle bundle.

Innervation and neurotrophic factors also impacted the dynamics of twitch contractions. In non-innervated muscle bundles without neurotrophic factors, the rise time (**Figure 4E**) and fall time (**Figure 4F**) decreased close to significantly and significantly, respectively, from one to two weeks of differentiation, which is consistent with a loss of slow fibers and/or gain of fast fibers. In contrast, bundles cultured with neurotrophic factors, with or without motor neurons, maintained more consistent rise and fall times from one to two weeks of differentiation (**Figure 4 E,F)**, possibly indicating less remodeling towards a fast, glycolytic phenotype.

### Impact of Innervation on Expression of Muscle Type Specification Genes

Motivated by the differences in rise and fall time observed between muscle bundles, we next measured the expression levels of a panel of hallmark fast and slow twitch genes using RT-qPCR. Because motor neurons were a relatively small component of innervated muscle tissues compared to muscle cells, we assumed they did not have a significant impact on the expression levels for the genes in our panel, which are also not expected to be expressed by neurons to an appreciable extent. At the one-week timepoint, non-innervated muscle bundles without neurotrophic factors expressed higher levels of the fast twitch gene myosin heavy chain (*MYH1A*) compared to muscle bundles with neurotrophic factors and motor neurons (**Figure 5A).** Slow myosin light chain 3 (*MYL3*) was also downregulated in non-innervated bundles with neurotrophic factors after one week compared to the other two conditions (**Figure 5A)**. All other genes showed similar levels of expression.

**Figure 5:**
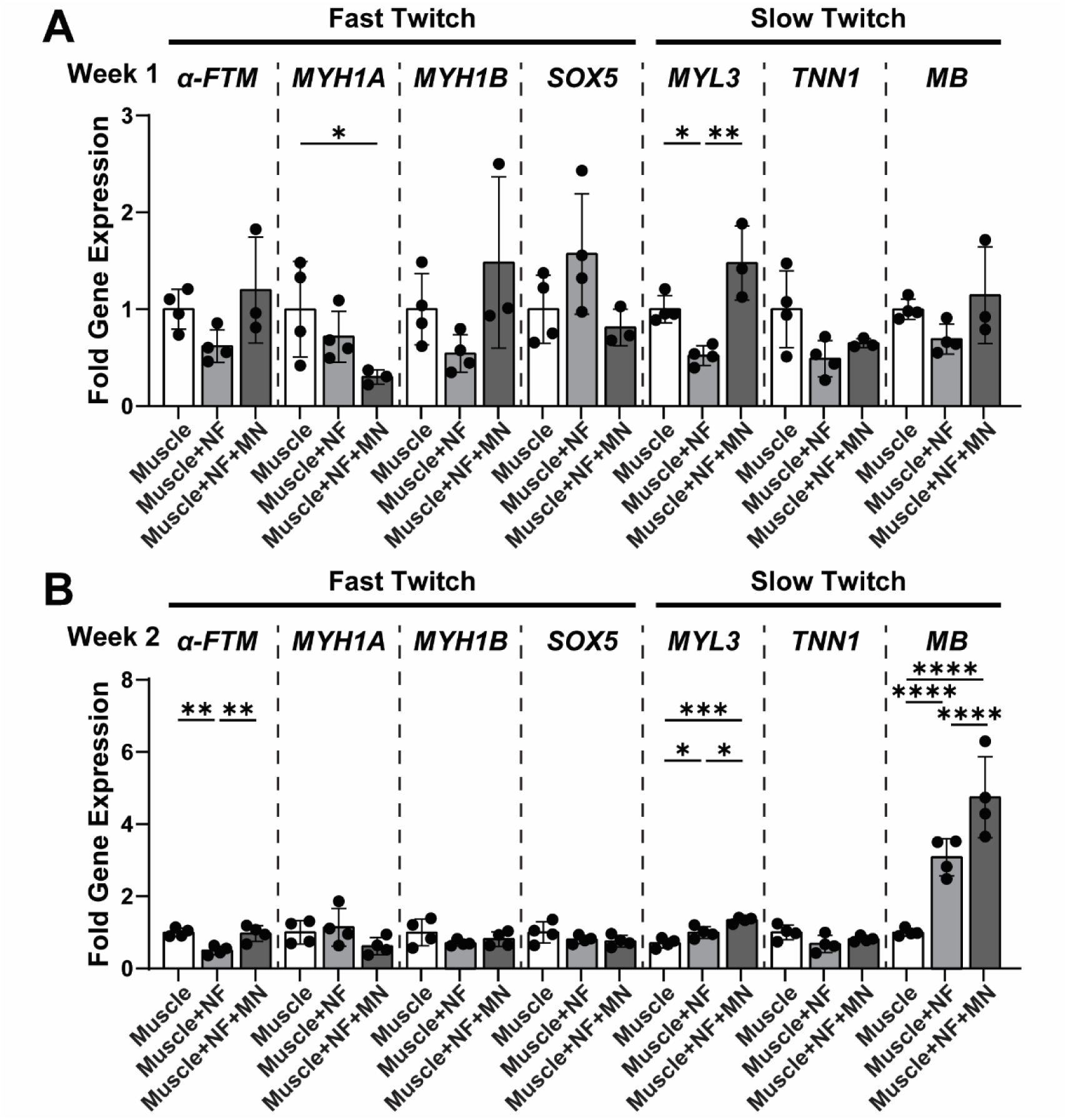
Expression of select fast and slow twitch muscle fiber genes in muscle bundles cultured with or without neurotrophic factors (NF) or motor neurons (MN). Fold change in gene expression after (A) one or (B) two weeks of culture for the following genes: α-tropomyosin (*α-FTM*), myosin heavy chain 1A (*MYH1A*), myosin heavy chain B (*MYHB*), sex-determining region Y-box 5 (*Sox5*), myosin light chain 3 (*MYL3*), troponin type 1 (*TNN1*), and myoglobin (*MB*).

After two weeks of differentiation, we observed a stark increase in the expression of myoglobin (*MB*) by innervated muscle bundles compared to non-innervated muscle bundles. Non-innervated muscle bundles with neurotrophic factors expressed *MB* to an intermediate level (**Figure 5B)**. Myoglobin is enriched in slow twitch muscle fibers because it stores and supplies oxygen and thus supports oxidative phosphorylation^31^. *MYL3* expression similarly increased stepwise from non-innervated bundles, to non-innervated bundles with neurotrophic factors, to innervated bundles (**Figure 5B).** *A-FTM*, encoding alpha-tropomyosin, a gene associated with fast twitch fibers^32^, was also downregulated in non-innervated bundles cultured with neurotrophic factors compared to the other two conditions (**Figure 5B).** Together, these data suggest that motor neuron innervation down-regulates *MYH1A* and up-regulates *MB* and *MYL3* in engineered skeletal muscle, consistent with a slow twitch phenotype. Neurotrophic factors alone also induce similar effects on gene expression, but to a lesser extent than the presence of both neurotrophic factors and motor neurons.

## Discussion

Motor neuron innervation is a critical regulator of skeletal muscle health and physiology *in vivo*^2^ and thus is also hypothesized to improve the physiology of engineered muscle constructs. Previous approaches to engineer innervated muscle tissue accomplished this by adding motor neurons to engineered muscle constructs post-differentiation, which resulted in a relatively minimal impact on the structural and functional maturity of the muscle^17, 19^. In this study, we mixed motor neurons and myoblasts together in a hydrogel solution, then proceeded to induce 3-D muscle differentiation, which led to a more pronounced improvement in the structure and contractile function of the muscle. Motor neuron innervation also enhanced the expression of genes related to oxidative phosphorylation and slow twitch fibers. Neurotrophic factors alone induced some changes in muscle phenotype, but to a lesser extent than innervation. These data reinforce the physiological importance of motor neurons on skeletal muscle development and maturation and suggest that motor neuron innervation could be further leveraged to engineer more mature muscle constructs for in vitro modeling or regenerative medicine.

Several engineered models of 2-D neuromuscular tissues have been successfully developed and implemented for mechanistic biology and neuromuscular disease modeling^10–13^. However, 3-D models more closely recapitulate the cell-cell and cell-matrix interactions of native tissues^26, 33,34^. Here, we fabricated chambers for engineering 3-D neuromuscular tissues using a benchtop, consumer-grade 3-D printer and laser cutter. Similar to other approaches that use benchtop fabrication equipment^14, 15, 17^, this approach is more accessible, cheaper, and faster than photolithography^16, 35^, enabling more rapid fabrication and prototyping. To measure force generation, we stimulated tissues with an external field stimulation electrode and then optically tracked the movement of a laser-cut, flexible PDMS rod. Similar optical tracking techniques are commonly used for measuring force generation in engineered muscle constructs^36, 37^. However, one limitation of optical tracking is that it is low-throughput and has relatively low sensitivity, which is a tradeoff of the simple fabrication method. Integration of stimulation electrodes and flexible electrical sensors into our device, as described in other systems^38–40^, would improve both the throughput and accuracy of our device.

Because we added motor neurons prior to muscle bundle compaction, we observed axons distributed throughout the volume of the bundle. By contrast, the axons of motor neurons added after muscle bundle formation likely struggle to migrate deep into the muscle and instead stay on the surface of the muscle^14, 16^. The increased interactions between muscle and neurons in our tissues may explain why we observed a greater effect of innervation on myogenic index and force generation compared to prior models. However, we were not able to confirm the presence of neuromuscular junctions in our tissues due to technical challenges in detecting these delicate structures. Previously, we did identify relatively mature neuromuscular junctions between these two cell types in 2-D tissue constructs^12^, providing some evidence that they likely form synapses in 3-D, but this would need to be confirmed. Alternatively, or additionally, the muscle tissue could be remodeling in response to paracrine crosstalk with motor neurons, as neurons can also secrete signaling factors, including BDNF and GDNF^41^. Establishing mechanisms of neuron-muscle crosstalk is an important topic for follow-up studies.

Innervation also impacted the dynamics of contraction and the expression of slow and fast twitch genes by our tissue constructs. Our starting muscle cells were isolated from the chick thigh muscle, which primarily comprises highly oxidative, slow muscle types^42^. Muscle bundles without neurotrophic factors or motor neurons experienced a significant decline in rise and fall time from one to two weeks, consistent with a transition from slow to fast fibers. By contrast, innervated muscle bundles and muscle bundles with neurotrophic factors maintained a more constant rise and fall time, suggesting that more slow fibers were maintained. These contractility results were also corroborated with RT-qPCR analysis, as innervated tissues expressed the lowest levels of *MYH1A* (fast twitch gene) at one week and the highest levels of *MYL3* and *MB* (slow twitch genes) at two weeks. *MYH1A* encodes for a myosin heavy chain protein expressed in fast type fibers and *MYL3* encodes for a myosin light chain protein expressed in slow type fibers. *MB* encodes for myoglobin, which is a selective marker for oxidative muscle fibers^43^. Myoglobin levels rapidly increase during neonatal development, as shown in mouse^44^. Continuous motor neuron stimulation also increases myoglobin levels *in vivo*, as shown in rabbit^44^, consistent with our *in vitro* results.

Electric pulse stimulation of non-innervated muscle tissues is commonly used to induce contraction and somewhat mimic innervation. Electrical pulse stimulation has been reported to enhance the expression of muscle genes, increase force generation, and induce a switch from fast to slow fiber types^45–47^. Electrical pulse stimulation also induces maturation in engineered 3-D cardiac tissues^35, 48^. Thus, it is plausible that motor neurons are improving the structure and function of our engineered muscle tissues by continuously stimulating muscle contraction, similar to electrical pulse stimulation. Although electrical pulse stimulation is a more straightforward approach, motor neuron innervation and electrical pulse stimulation have been shown to induce distinct phenotypes in engineered muscle tissues^49, 50^. Thus, electrical pulse stimulation is not completely equivalent to motor neuron innervation and may induce some non-natural phenotypes that are undesirable. Furthermore, other forms of paracrine interactions between muscle and neurons are absent with electrical pulse stimulation, which likely also impact tissue phenotypes. Paracrine crosstalk between motor neurons and skeletal muscle involves a complex mixture of neurotrophic factors and other myokines not investigated here^51, 52^, which may further explain the higher impact of motor neuron co-culture compared to neurotrophic factors alone. Future studies can focus on determining the precise mechanisms of muscle enhancement induced by motor neurons.

In this study, we co-cultured motor neurons within engineered muscle bundles and observed enhanced muscle maturation and maintenance of slow twitch phenotypes. However, there are several caveats and limitations of our study that should be considered. First, we used human iPSC-derived motor neurons and primary embryonic chick muscle cells. We chose chick cells because they are highly accessible in large quantities and can quickly and easily differentiate into highly contractile muscle tissues^12^. Previous work has also shown synapse formation between human iPSC-derived motor neurons and chick muscle^12, 21, 53^. However, the use of chick cells does significantly limit human relevance. Thus, future studies should focus on using primary human myoblasts or human iPSC-derived myoblasts^54^ to better model human neuromuscular development. Another limitation is that we did not directly confirm or evaluate neuromuscular junction structure or function. Thus, the mechanisms of how innervation improves muscle maturation is an important topic for future study. In the reciprocal direction, there may also be beneficial effects of muscle on the motor neurons that were not explored in this study. We also neglected many other important cell types that are present in endogenous muscle, such as fibroblasts^30^, endothelial cells^55^, and macrophages^33^, which have also been co-cultured with engineered muscle constructs and shown to improve muscle phenotype. Thus, cell-cell interactions with supporting cell types, including, but not limited to, motor neurons, are key regulators of skeletal muscle development and maturation and are an important design feature to consider for ongoing efforts in engineering mature skeletal muscle tissue constructs for disease modeling or transplantation.

## Acknowledgements

We thank the NINDS Biorepository at Coriell Institute for providing human cell lines used for this study. This work was funded by NSF GRFP grant DGE1418060 (JWS); USC Broad Innovation Award (JKI, MLM); Broad Collaborative Challenge Grants for Graduate Students and Postdoctoral Fellows (JWS); USC Viterbi Internal Center Incubator and Center for Integrated Electronic and Biological Organisms (CIEBOrg) (MLM); NSF CAREER Award 1944734 (MLM); CIRM Research Training Grant EDUC4-12756 (SKD); grant 2023-332386 from the Chan Zuckerberg Initiative DAF, an advised fund of Silicon Valley Community Foundation (MLM); ARCS Los Angeles Founder Chapter (SKD); NIH grants R01NS097850 (JKI), 2R01NS097850-05 (JKI), and 3R01NS097850-01S1 (EH); US Department of Defense grants W81XWH-15-1-0187 and W81XWH-20-1-0424 (JKI). Additional funding was provided from grants from the Tau Consortium (JKI), the New York Stem Cell Foundation (JKI), the John Douglas French Alzheimer′s Foundation (JKI), and the Merkin Family Foundation (JKI). JKI is a New York Stem Cell Foundation-Robertson Investigator and the John Douglas French Alzheimer’s Endowed Professor.

## References

(1) Harris, A. J. Embryonic growth and innervation of rat skeletal muscles. I. Neural regulation of muscle fibre numbers. Philos Trans R Soc Lond B Biol Sci 1981, 293 (1065), 257–277. DOI: 10.1098/rstb.1981.0076 From NLM Medline.

(2) Tintignac, L. A.; Brenner, H. R.; Ruegg, M. A. Mechanisms Regulating Neuromuscular Junction Development and Function and Causes of Muscle Wasting. Physiol Rev 2015, 95 (3), 809–852. DOI: 10.1152/physrev.00033.2014 From NLM Medline.

(3) Mu, L.; Sobotka, S.; Su, H. Nerve-muscle-endplate band grafting: a new technique for muscle reinnervation. Neurosurgery 2011, 69 (2 Suppl Operative), ons208–224; discussion ons224. DOI: 10.1227/NEU.0b013e31822ed596 From NLM Medline.

(4) Whalen, R. G.; Harris, J. B.; Butler-Browne, G. S.; Sesodia, S. Expression of myosin isoforms during notexin-induced regeneration of rat soleus muscles. Dev Biol 1990, 141 (1), 24–40. DOI: 10.1016/0012-1606(90)90099-5 From NLM Medline.

(5) Lauretani, F.; Bandinelli, S.; Bartali, B.; Di Iorio, A.; Giacomini, V.; Corsi, A. M.; Guralnik, J. M.; Ferrucci, L. Axonal degeneration affects muscle density in older men and women. Neurobiol Aging 2006, 27 (8), 1145–1154. DOI: 10.1016/j.neurobiolaging.2005.06.009 From NLM Medline.

(6) Day, J. W.; Howell, K.; Place, A.; Long, K.; Rossello, J.; Kertesz, N.; Nomikos, G. Advances and limitations for the treatment of spinal muscular atrophy. BMC Pediatr 2022, 22 (1), 632. DOI: 10.1186/s12887-022-03671-x From NLM Medline.

(7) Ling, K. K.; Gibbs, R. M.; Feng, Z.; Ko, C. P. Severe neuromuscular denervation of clinically relevant muscles in a mouse model of spinal muscular atrophy. Hum Mol Genet 2012, 21 (1), 185–195. DOI: 10.1093/hmg/ddr453 From NLM Medline.

(8) Chen, C. J.; Cheng, F. C.; Su, H. L.; Sheu, M. L.; Lu, Z. H.; Chiang, C. Y.; Yang, D. Y.; Sheehan, J.; Pan, H. C. Improved neurological outcome by intramuscular injection of human amniotic fluid derived stem cells in a muscle denervation model. PLoS One 2015, 10 (5), e0124624. DOI: 10.1371/journal.pone.0124624 From NLM Medline.

(9) Helgren, M. E.; Squinto, S. P.; Davis, H. L.; Parry, D. J.; Boulton, T. G.; Heck, C. S.; Zhu, Y.; Yancopoulos, G. D.; Lindsay, R. M.; DiStefano, P. S. Trophic effect of ciliary neurotrophic factor on denervated skeletal muscle. Cell 1994, 76 (3), 493–504. DOI: 10.1016/0092-8674(94)90113-9 From NLM Medline.

(10) Happe, C. L.; Tenerelli, K. P.; Gromova, A. K.; Kolb, F.; Engler, A. J. Mechanically patterned neuromuscular junctions-in-a-dish have improved functional maturation. Mol Biol Cell 2017, 28 (14), 1950–1958. DOI: 10.1091/mbc.E17-01-0046 From NLM Medline.

(11) Santhanam, N.; Kumanchik, L.; Guo, X.; Sommerhage, F.; Cai, Y.; Jackson, M.; Martin, C.; Saad, G.; McAleer, C. W.; Wang, Y.;, et al. Stem cell derived phenotypic human neuromuscular junction model for dose response evaluation of therapeutics. Biomaterials 2018, 166, 64–78. DOI: 10.1016/j.biomaterials.2018.02.047 From NLM Medline.

(12) Santoso, J. W.; Li, X.; Gupta, D.; Suh, G. C.; Hendricks, E.; Lin, S.; Perry, S.; Ichida, J. K.; Dickman, D.; McCain, M. L. Engineering skeletal muscle tissues with advanced maturity improves synapse formation with human induced pluripotent stem cell-derived motor neurons. APL Bioeng 2021, 5 (3), 036101. DOI: 10.1063/5.0054984 From NLM PubMed-not-MEDLINE.

(13) Steinbeck, J. A.; Jaiswal, M. K.; Calder, E. L.; Kishinevsky, S.; Weishaupt, A.; Toyka, K. V.; Goldstein, P. A.; Studer, L. Functional Connectivity under Optogenetic Control Allows Modeling of Human Neuromuscular Disease. Cell Stem Cell 2016, 18 (1), 134–143. DOI: 10.1016/j.stem.2015.10.002 From NLM Medline.

(14) Afshar Bakooshli, M.; Lippmann, E. S.; Mulcahy, B.; Iyer, N.; Nguyen, C. T.; Tung, K.; Stewart, B. A.; van den Dorpel, H.; Fuehrmann, T.; Shoichet, M.;, et al. A 3D culture model of innervated human skeletal muscle enables studies of the adult neuromuscular junction. Elife 2019, 8. DOI: 10.7554/eLife.44530 From NLM Medline.

(15) Morimoto, Y.; Kato-Negishi, M.; Onoe, H.; Takeuchi, S. Three-dimensional neuron-muscle constructs with neuromuscular junctions. Biomaterials 2013, 34 (37), 9413–9419. DOI: 10.1016/j.biomaterials.2013.08.062 From NLM Medline.

(16) Osaki, T.; Uzel, S. G. M.; Kamm, R. D. On-chip 3D neuromuscular model for drug screening and precision medicine in neuromuscular disease. Nat Protoc 2020, 15 (2), 421–449. DOI: 10.1038/s41596-019-0248-1 From NLM Medline.

(17) Rimington, R. P.; Fleming, J. W.; Capel, A. J.; Wheeler, P. C.; Lewis, M. P. Bioengineered model of the human motor unit with physiologically functional neuromuscular junctions. Sci Rep 2021, 11 (1), 11695. DOI: 10.1038/s41598-021-91203-5 From NLM Medline.

(18) Santoso, J. W.; McCain, M. L. Neuromuscular disease modeling on a chip. Dis Model Mech 2020, 13 (7). DOI: 10.1242/dmm.044867 From NLM Medline.

(19) Smith, A. S.; Passey, S. L.; Martin, N. R.; Player, D. J.; Mudera, V.; Greensmith, L.; Lewis, M. P. Creating Interactions between Tissue-Engineered Skeletal Muscle and the Peripheral Nervous System. Cells Tissues Organs 2016, 202 (3-4), 143–158. DOI: 10.1159/000443634 From NLM Medline.

(20) Ariyasinghe, N. R.; Santoso, J. W.; Gupta, D.; Pincus, M. J.; August, P. R.; McCain, M. L. Optical Clearing of Skeletal Muscle Bundles Engineered in 3-D Printed Templates. Ann Biomed Eng 2021, 49 (2), 523–535. DOI: 10.1007/s10439-020-02583-0 From NLM Medline.

(21) Son, E. Y.; Ichida, J. K.; Wainger, B. J.; Toma, J. S.; Rafuse, V. F.; Woolf, C. J.; Eggan, K. Conversion of mouse and human fibroblasts into functional spinal motor neurons. Cell Stem Cell 2011, 9 (3), 205–218. DOI: 10.1016/j.stem.2011.07.014 From NLM Medline.

(22) Shi, Y.; Lin, S.; Staats, K. A.; Li, Y.; Chang, W. H.; Hung, S. T.; Hendricks, E.; Linares, G. R.; Wang, Y.; Son, E. Y.;, et al. Haploinsufficiency leads to neurodegeneration in C9ORF72 ALS/FTD human induced motor neurons. Nat Med 2018, 24 (3), 313–325. DOI: 10.1038/nm.4490 From NLM Medline.

(23) Okita, K.; Matsumura, Y.; Sato, Y.; Okada, A.; Morizane, A.; Okamoto, S.; Hong, H.; Nakagawa, M.; Tanabe, K.; Tezuka, K.;, et al. A more efficient method to generate integration-free human iPS cells. Nat Methods 2011, 8 (5), 409–412. DOI: 10.1038/nmeth.1591 From NLM Medline.

(24) Du, Z. W.; Chen, H.; Liu, H.; Lu, J.; Qian, K.; Huang, C. L.; Zhong, X.; Fan, F.; Zhang, S. C. Generation and expansion of highly pure motor neuron progenitors from human pluripotent stem cells. Nat Commun 2015, 6, 6626. DOI: 10.1038/ncomms7626 From NLM Medline.

(25) Davis, B. N. J.; Santoso, J. W.; Walker, M. J.; Oliver, C. E.; Cunningham, M. M.; Boehm, C. A.; Dawes, D.; Lasater, S. L.; Huffman, K.; Kraus, W. E.;, et al. Modeling the Effect of TNF-alpha upon Drug-Induced Toxicity in Human, Tissue-Engineered Myobundles. Ann Biomed Eng 2019, 47 (7), 1596–1610. DOI: 10.1007/s10439-019-02263-8 From NLM Medline.

(26) Madden, L.; Juhas, M.; Kraus, W. E.; Truskey, G. A.; Bursac, N. Bioengineered human myobundles mimic clinical responses of skeletal muscle to drugs. Elife 2015, 4, e04885. DOI: 10.7554/eLife.04885 From NLM Medline.

(27) Juhas, M.; Bursac, N. Roles of adherent myogenic cells and dynamic culture in engineered muscle function and maintenance of satellite cells. Biomaterials 2014, 35 (35), 9438–9446. DOI: 10.1016/j.biomaterials.2014.07.035 From NLM Medline.

(28) Livak, K. J.; Schmittgen, T. D. Analysis of relative gene expression data using real-time quantitative PCR and the 2(-Delta Delta C(T)) Method. Methods 2001, 25 (4), 402–408. DOI: 10.1006/meth.2001.1262 From NLM Medline.

(29) Delezie, J.; Weihrauch, M.; Maier, G.; Tejero, R.; Ham, D. J.; Gill, J. F.; Karrer-Cardel, B.; Ruegg, M. A.; Tabares, L.; Handschin, C. BDNF is a mediator of glycolytic fiber-type specification in mouse skeletal muscle. Proc Natl Acad Sci U S A 2019, 116 (32), 16111–16120. DOI: 10.1073/pnas.1900544116 From NLM Medline.

(30) Li, M.; Dickinson, C. E.; Finkelstein, E. B.; Neville, C. M.; Sundback, C. A. The role of fibroblasts in self-assembled skeletal muscle. Tissue Eng Part A 2011, 17 (21-22), 2641–2650. DOI: 10.1089/ten.TEA.2010.0700 From NLM Medline.

(31) Okumura, N.; Hashida-Okumura, A.; Kita, K.; Matsubae, M.; Matsubara, T.; Takao, T.; Nagai, K. Proteomic analysis of slow- and fast-twitch skeletal muscles. Proteomics 2005, 5 (11), 2896–2906. DOI: 10.1002/pmic.200401181 From NLM Medline.

(32) Lemonnier, M.; Balvay, L.; Mouly, V.; Libri, D.; Fiszman, M. Y. The chicken gene encoding the alpha isoform of tropomyosin of fast-twitch muscle fibers: organization, expression and identification of the major proteins synthesized. Gene 1991, 107 (2), 229–240. DOI: 10.1016/0378-1119(91)90323-4 From NLM Medline.

(33) Juhas, M.; Abutaleb, N.; Wang, J. T.; Ye, J.; Shaikh, Z.; Sriworarat, C.; Qian, Y.; Bursac, N. Incorporation of macrophages into engineered skeletal muscle enables enhanced muscle regeneration. Nat Biomed Eng 2018, 2 (12), 942–954. DOI: 10.1038/s41551-018-0290-2 From NLM PubMed-not-MEDLINE.

(34) Khodabukus, A. Tissue-Engineered Skeletal Muscle Models to Study Muscle Function, Plasticity, and Disease. Front Physiol 2021, 12, 619710. DOI: 10.3389/fphys.2021.619710 From NLM PubMed-not-MEDLINE.

(35) Nunes, S. S.; Miklas, J. W.; Liu, J.; Aschar-Sobbi, R.; Xiao, Y.; Zhang, B.; Jiang, J.; Masse, S.; Gagliardi, M.; Hsieh, A.;, et al. Biowire: a platform for maturation of human pluripotent stem cell-derived cardiomyocytes. Nat Methods 2013, 10 (8), 781–787. DOI: 10.1038/nmeth.2524 From NLM Medline.

(36) Bliley, J. M.; Vermeer, M.; Duffy, R. M.; Batalov, I.; Kramer, D.; Tashman, J. W.; Shiwarski, D. J.; Lee, A.; Teplenin, A. S.; Volkers, L.;, et al. Dynamic loading of human engineered heart tissue enhances contractile function and drives a desmosome-linked disease phenotype. Sci Transl Med 2021, 13 (603). DOI: 10.1126/scitranslmed.abd1817 From NLM Medline.

(37) Qu, Y.; Feric, N.; Pallotta, I.; Singh, R.; Sobbi, R.; Vargas, H. M. Inotropic assessment in engineered 3D cardiac tissues using human induced pluripotent stem cell-derived cardiomyocytes in the Biowire(TM) II platform. J Pharmacol Toxicol Methods 2020, 105, 106886. DOI: 10.1016/j.vascn.2020.106886 From NLM Medline.

(38) Lind, J. U.; Busbee, T. A.; Valentine, A. D.; Pasqualini, F. S.; Yuan, H.; Yadid, M.; Park, S. J.; Kotikian, A.; Nesmith, A. P.; Campbell, P. H.;, et al. Instrumented cardiac microphysiological devices via multimaterial three-dimensional printing. Nat Mater 2017, 16 (3), 303–308. DOI: 10.1038/nmat4782 From NLM Medline.

(39) Wu, Q.; Zhang, P.; O’Leary, G.; Zhao, Y.; Xu, Y.; Rafatian, N.; Okhovatian, S.; Landau, S.; Valiante, T. A.; Travas-Sejdic, J.;, et al. Flexible 3D printed microwires and 3D microelectrodes for heart-on-a-chip engineering. Biofabrication 2023, 15 (3). DOI: 10.1088/1758-5090/acd8f4 From NLM Medline.

(40) Yip, J. K.; Sarkar, D.; Petersen, A. P.; Gipson, J. N.; Tao, J.; Kale, S.; Rexius-Hall, M. L.; Cho, N.; Khalil, N. N.; Kapadia, R.;, et al. Contact photolithography-free integration of patterned and semi-transparent indium tin oxide stimulation electrodes into polydimethylsiloxane-based heart-on-a-chip devices for streamlining physiological recordings. Lab Chip 2021, 21 (4), 674–687. DOI: 10.1039/d0lc00948b From NLM Medline.

(41) Deng, J.; Ji, Y.; Zhu, F.; Liu, L.; Li, L.; Bai, X.; Li, H.; Liu, X.; Luo, Y.; Lin, B.;, et al. Mapping secretome-mediated interaction between paired neuron-macrophage single cells. Proc Natl Acad Sci U S A 2022, 119 (44), e2200944119. DOI: 10.1073/pnas.2200944119 From NLM Medline.

(42) Weng, K.; Huo, W.; Li, Y.; Zhang, Y.; Zhang, Y.; Chen, G.; Xu, Q. Fiber characteristics and meat quality of different muscular tissues from slow- and fast-growing broilers. Poult Sci 2022, 101 (1), 101537. DOI: 10.1016/j.psj.2021.101537 From NLM Medline.

(43) Bekedam, M. A.; van Beek-Harmsen, B. J.; van Mechelen, W.; Boonstra, A.; van der Laarse, W. J. Myoglobin concentration in skeletal muscle fibers of chronic heart failure patients. J Appl Physiol (1985) 2009, 107 (4), 1138–1143. DOI: 10.1152/japplphysiol.00149.2009 From NLM Medline.

(44) Garry, D. J.; Bassel-Duby, R. S.; Richardson, J. A.; Grayson, J.; Neufer, P. D.; Williams, R. S. Postnatal development and plasticity of specialized muscle fiber characteristics in the hindlimb. Dev Genet 1996, 19 (2), 146–156. DOI: 10.1002/(SICI)1520-6408(1996)19:2<146::AID-DVG6>3.0.CO;2-9 From NLM Medline.

(45) Charoensook, S. N.; Williams, D. J.; Chakraborty, S.; Leong, K. W.; Vunjak-Novakovic, G. Bioreactor model of neuromuscular junction with electrical stimulation for pharmacological potency testing. Integr Biol (Camb) 2017, 9 (12), 956–967. DOI: 10.1039/c7ib00144d From NLM Medline.

(46) Ito, A.; Yamamoto, Y.; Sato, M.; Ikeda, K.; Yamamoto, M.; Fujita, H.; Nagamori, E.; Kawabe, Y.; Kamihira, M. Induction of functional tissue-engineered skeletal muscle constructs by defined electrical stimulation. Sci Rep 2014, 4, 4781. DOI: 10.1038/srep04781 From NLM Medline.

(47) Nedachi, T.; Fujita, H.; Kanzaki, M. Contractile C2C12 myotube model for studying exercise-inducible responses in skeletal muscle. Am J Physiol Endocrinol Metab 2008, 295 (5), E1191–1204. DOI: 10.1152/ajpendo.90280.2008 From NLM Medline.

(48) Ronaldson-Bouchard, K.; Ma, S. P.; Yeager, K.; Chen, T.; Song, L.; Sirabella, D.; Morikawa, K.; Teles, D.; Yazawa, M.; Vunjak-Novakovic, G. Advanced maturation of human cardiac tissue grown from pluripotent stem cells. Nature 2018, 556 (7700), 239–243. DOI: 10.1038/s41586-018-0016-3 From NLM Medline.

(49) Jan, V.; Mis, K.; Nikolic, N.; Dolinar, K.; Petric, M.; Bone, A.; Thoresen, G. H.; Rustan, A. C.; Mars, T.; Chibalin, A. V.;, et al. Effect of differentiation, de novo innervation, and electrical pulse stimulation on mRNA and protein expression of Na+,K+-ATPase, FXYD1, and FXYD5 in cultured human skeletal muscle cells. PLoS One 2021, 16 (2), e0247377. DOI: 10.1371/journal.pone.0247377 From NLM Medline.

(50) Mars, T.; Mis, K.; Meznaric, M.; Prpar Mihevc, S.; Jan, V.; Haugen, F.; Rogelj, B.; Rustan, A. C.; Thoresen, G. H.; Pirkmajer, S.;, et al. Innervation and electrical pulse stimulation - in vitro effects on human skeletal muscle cells. Appl Physiol Nutr Metab 2021, 46 (4), 299–308. DOI: 10.1139/apnm-2019-0575 From NLM Medline.

(51) Saini, J.; Faroni, A.; Reid, A. J.; Mouly, V.; Butler-Browne, G.; Lightfoot, A. P.; McPhee, J. S.; Degens, H.; Al-Shanti, N. Cross-talk between motor neurons and myotubes via endogenously secreted neural and muscular growth factors. Physiol Rep 2021, 9 (8), e14791. DOI: 10.14814/phy2.14791 From NLM Medline.

(52) Huang, K. Y.; Upadhyay, G.; Ahn, Y.; Sakakura, M.; Pagan-Diaz, G. J.; Cho, Y.; Weiss, A. C.; Huang, C.; Mitchell, J. W.; Li, J.;, et al. Neuronal innervation regulates the secretion of neurotrophic myokines and exosomes from skeletal muscle. Proc Natl Acad Sci U S A 2024, 121 (19), e2313590121. DOI: 10.1073/pnas.2313590121 From NLM Medline.

(53) Toma, J. S.; Shettar, B. C.; Chipman, P. H.; Pinto, D. M.; Borowska, J. P.; Ichida, J. K.; Fawcett, J. P.; Zhang, Y.; Eggan, K.; Rafuse, V. F. Motoneurons derived from induced pluripotent stem cells develop mature phenotypes typical of endogenous spinal motoneurons. J Neurosci 2015, 35 (3), 1291–1306. DOI: 10.1523/JNEUROSCI.2126-14.2015 From NLM Medline.

(54) Pinton, L.; Khedr, M.; Lionello, V. M.; Sarcar, S.; Maffioletti, S. M.; Dastidar, S.; Negroni, E.; Choi, S.; Khokhar, N.; Bigot, A.;, et al. 3D human induced pluripotent stem cell-derived bioengineered skeletal muscles for tissue, disease and therapy modeling. Nat Protoc 2023, 18 (4), 1337–1376. DOI: 10.1038/s41596-022-00790-8 From NLM Medline.

(55) Osaki, T.; Sivathanu, V.; Kamm, R. D. Crosstalk between developing vasculature and optogenetically engineered skeletal muscle improves muscle contraction and angiogenesis. Biomaterials 2018, 156, 65–76. DOI: 10.1016/j.biomaterials.2017.11.041 From NLM Medline.

