## Supplemental Information for "Human iPSC-derived motor neuron innervation enhances the differentiation of muscle bundles engineered with benchtop fabrication techniques"

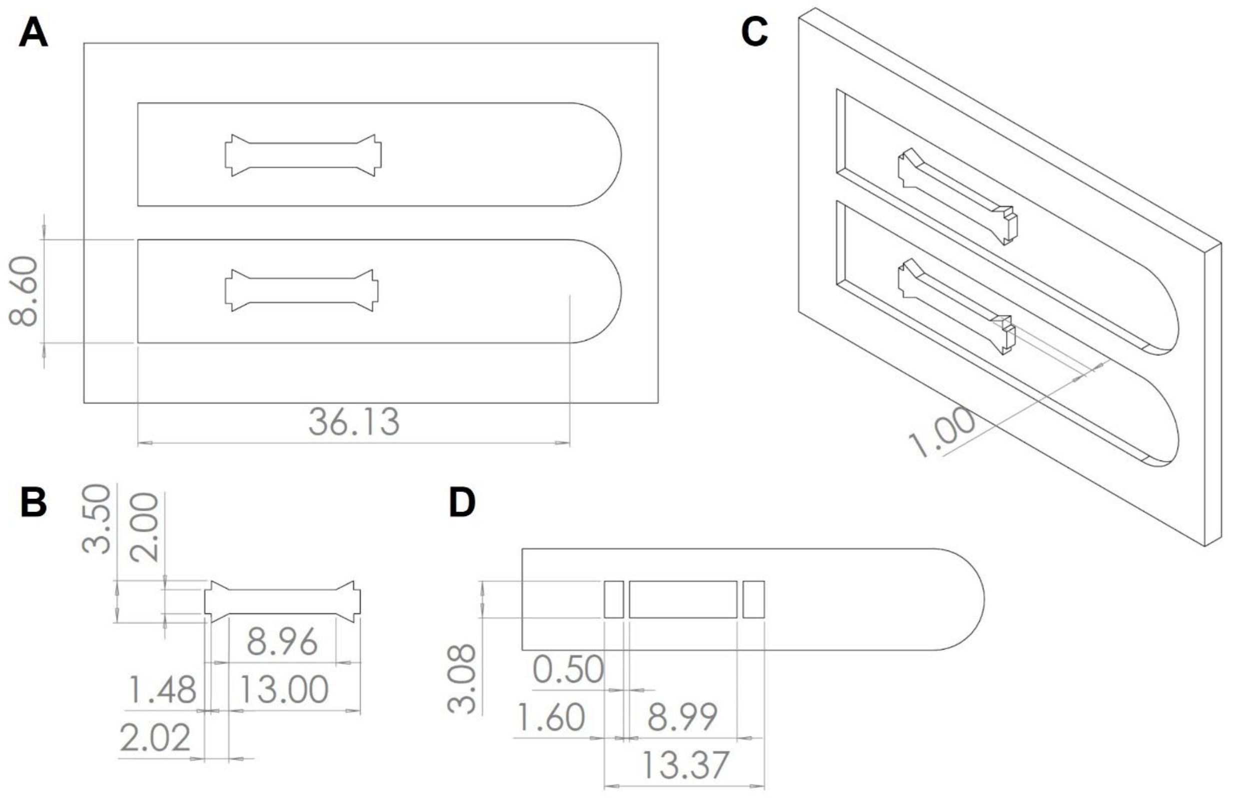


**Supplemental Figure 1: Design details for muscle bundle devices.** (A) Top view of 3-D printed template comprising two half-chambers. Round tabs enable easy removal of PDMS from printed templates. (B) Muscle bundle chamber shown in more detail. (C) 3-D view of template that was 3-D printed and then cast with PDMS to form the top and bottom of muscle bundle devices. (D) Top view of anchor rod design that was laser cut into silicone sheet and bonded between the top and bottom pieces described in (C).

| Gene Symbol | Forward/Reverse | Anneal temperature (°C) | Product size | Type |
|---|---|---|---|---|
| Alpha FTM | F: CCTGCTCTGATTTCGGCTGT<br>R:TCTTGATGGCATCCATGGCG | 60 | 71 bp | Fast twitch |
| MYH1A | F: TCTTCCAGTCAGCACAAGACCT<br>R: GACTTTCGGAGGTAGGGAGCG | 61 | 134 bp | Fast twitch |
| MYH1B | F: AAGTTCCGCAAGATCCAGCA<br>R: ATGCAGAGGAATCTATGGTCTTT | 60 | 257 bp | Fast twitch |
| SOX5 | F: TTAACGCGGGGAGTTAGACG<br>R: GACAAAGCTTTTCCCCTGCG | 60 | 90 bp | Fast twitch |
| MYL3 | F: GCACTGGTGACTTTTCCTGC<br>R: GTTCAGGCGCCTTCTTAGGT | 60 | 120 bp | Slow twitch |
| TNNI1 | F: CGGAGCCCAGGGAGAGAAAA<br>R:CTGCTCCCACTCCTCCTTGG | 61 | 95 bp | Slow twitch |
| MB | F:CAACCGCCATAGTCAGCACT<br>R: AGGACTTGTTGCCACTCCTG | 60 | 91 bp | Slow twitch |
| ACTB | F: CGGACTGTTACCAACACCCA<br>R: CCTGAGTCAAGCGCCAAAAG | 60 | 114 bp | Housekeeping |

**Supplemental Table 1: Primer sequences for qPCR.**

**Supplemental Video 1:** Contracting innervated muscle bundle. Muscle bundle with hiPSC-derived motor neurons at two-week timepoint. Stimulation was applied at 2 Hz (twitch) followed by 20 Hz (tetanus).

**Supplemental Video 2:** Contracting non-innervated muscle bundle. Muscle bundle without hiPSC-derived motor neurons at two-week timepoint. Stimulation was applied at 2 Hz (twitch) followed by 20 Hz (tetanus).
